# FBXW7β isoform drives transcriptional activation of a proinflammatory TNF cluster in normal and malignant pro-B cells

**DOI:** 10.1101/2022.04.24.489313

**Authors:** Scarlett Y. Yang, Katharina E. Hayer, Hossein Fazelinia, Lynn A. Spruce, Mukta Asnani, Kathryn L. Black, Ammar S. Naqvi, Vinodh Pillai, Yoseph Barash, Kojo S. J. Elenitoba-Johnson, Andrei Thomas-Tikhonenko

## Abstract

Non-canonical exon usage plays many important roles in cellular phenotypes, but its contribution to human B-cell development remains sketchily understood. To fill this gap, we collected various B-cell fractions from bone marrow and tonsil donors, performed RNA-seq, and examined transcript variants. We identified 150 genes that harbor local splicing variations in all pairwise comparisons. One of them encodes FBXW7, an E3 ubiquitin ligase implicated as a cancer driver in several blood cancers. Surprisingly, we discovered that in normal human pro-B cells, the predominant transcript utilized an alternative first exon to produce the poorly characterized FBXW7β isoform, previously thought to be restricted to neural tissues. The FBXW7β transcript was also abundant in cell lines and primary samples of pediatric B-cell acute lymphoblastic leukemia (B-ALL), which originates in the bone marrow. When overexpressed in a heterologous cell system, this transcript yielded the expected protein product, as judged by anti-FLAG immunoblotting and mass spectrometry. Furthermore, in REH B-ALL cells, FBXW7β mRNA was the only FBXW7 isoform enriched in the polyribosome fraction. To shed light on possible functions of FBXW7β, we utilized gain- and loss-of-function approaches and identified an FBXW7β-dependent inflammatory gene signature, apparent in a subset of B-ALL with high FBXW7β expression. This signature contained several members of the TNF superfamily, including those comprising the HLA Class III cluster (LTB, LST1, NCR3, LTA, and NFKBIL1). Our findings suggest that FBXW7β expression drives proinflammatory responses, which could contribute to normal B-cell development, leukemogenesis and responses to anti-cancer therapies.

**Key points:**

- Previously thought to be restricted to neural tissues, FBXW7β is the predominant FBXW7 isoform in normal and malignant human pro-B cells.
- FBXW7β promotes transcriptional activation of a proinflammatory gene cluster that contains TNF superfamily members.

## Introduction

During normal B-cell development and neoplastic transformation, cellular transcriptomes undergo dramatic changes. While these changes can be regulated by transcriptional activation and repression, transcript variation is another major mechanism that allows versatile gene regulation in response to cellular and environmental cues. Through tissue- or developmental stage-specific expression of distinct RNA isoforms from one given gene, transcript variation increases proteome diversity.

Transcript variation occurs via multiple contributing mechanisms, which could be divided into three major categories: alternative splicing (AS), alternative polyadenylation (APA), and alternative first exon usage (AFE). AS relies on the spliceosome for exon inclusion or exclusion. In contrast, APA and AFE are not directly mediated by the spliceosome, therefore considered non-AS events.^1^ While APA involves different poly(A) site selection for pre-mRNA cleavage and the addition of a poly(A) tail, AFE isoforms are driven by distinct promoters. Nevertheless, regulation of these three types of transcript variation appears interrelated.^1^

There is a growing appreciation of the importance of transcript variation in immune system development.^2,3^ For example, the two splicing variants of the E2A transcription factor regulate early B-cell differentiation and class switching.^4^ However, the extent to which transcript variation contributes to B-cell development and malignancies is not clear, especially in humans. To this end, we previously examined splicing events in B-cell acute lymphoblastic leukemia (B-ALL) and found widespread aberrant splicing.^5^ Most of the top 20 mutated genes in B-ALL also exhibit transcript variation.^5^ Interestingly, among B-ALL samples, the prevalence of transcript variation far exceeds that of somatic mutations in these genes.^5^ One salient example is FBXW7, which is included in the Catalog of Somatic Mutations in Cancer (COSMIC) as a curated Tier 1 Census gene and Cancer Hallmark gene.^6^

FBXW7 has been suggested to play different roles in cancer. It can act as a tumor suppressor or an oncogene in a context-dependent manner.^7–10^ To add to the complexity, alternative first exons of *FBXW7* give rise to three coding isoforms: α, γ, and β. The protein isoforms localize to distinct cellular compartments and exhibit different tissue distribution.^11,12^ The α isoform is ubiquitously expressed and extensively studied. In contrast, the expression and function of the β isoform is not fully characterized. While Fbxw7β is expressed in mouse brains^13^, to our knowledge, there is no evidence of Fbxw7β expression in hematopoietic tissues in mice. Expression of the β isoform has been reported in normal human brain, breast, testis, and assorted cancer tissues.^14–18^ However, due to technical constraints, such studies were often limited to RNA instead of protein analysis. Additionally, *FBXW7β* mRNA structure was inferred from individual exon junctions, lacking evidence for the corresponding full-length transcripts. Finally, while experiments with epitope-tagged FBXW7β protein offered interesting insights into its subcellular localization^19^, research on its endogenous counterpart was curbed by the lack of isoform-specific antibodies. As a result, relative contributions of the various FBXW7 isoforms to normal development and malignant transformation, especially in the hematological system, remains poorly understood.

To elucidate the role of FBXW7 isoforms in leukemia and normal human B-cell development, we employed deep short-read RNA-seq to discern isoform expression in normal progenitor subsets and primary B-cell Acute Lymphoblastic Leukemia (B-ALL) patient samples. We found that *FBXW7β* is the predominant transcript isoform in normal human pro-B cells, while α is the predominant isoform at other stages of development. We detected full-length *FBXW7β* transcript expression in B-ALL cells using long-read RNA sequencing and demonstrated its translatability using mass spectrometry and polysome profiling. With isoform-specific knockdown, knockout, and re-constitution systems, we show that FBXW7β regulates the TNF superfamily signaling pathways in B-ALL cells. Such regulation underpins the potential role of FBXW7β and its downstream effectors in leukemogenesis and anti-tumor cytotoxicity.

## Methods

*Additional information is contained in the Supplemental Methods*.

### Primary sample acquisition from Children’s Hospital of Philadelphia (CHOP)

According to 45 CFR 102(f), research on deidentified, discarded, or excess specimens does not constitute human subject research. IRB oversight was therefore not required for analyzing the normal pediatric bone marrow and tonsil samples. Eighteen primary pediatric B-ALL samples were obtained from the CHOP Center for Childhood Cancer Research leukemia biorepository as previously described.^5^

### Bone marrow fractionation

Normal human mononuclear cells and whole bone marrow aspirates underwent Ficoll gradient and Fluorescence-Activated Cell Sorting (FACS) selection as previously described.^5^ Live cells were sorted into early progenitor (CD34^+^CD19^-^IgM^-^), pro-B (CD34^+^CD19^+^IgM^-^), pre-B (CD34^-^CD19^+^IgM^-^), and immature B cell (CD34^-^CD19^+^IgM^+^) fractions.

### Naïve mature B cell isolation

Normal human tonsils from CHOP were manually homogenized with Precision Large Tissue Grinder (Covidien 3500SA) and passed through a 70 μm strainer. Cells were washed with PBS, blocked with Human TruStain (Biolegend), and stained with anti-CD19 APC, anti-IgM FITC, anti-IgD PE. Cells were pelleted at 250 × g for 5 min at 4°C and washed twice in PBS. Cells were resuspended in 2% FBS and 0.1 μg/mL DAPI in PBS. CD19^+^IgM^+^IgD^+^ Cells were sorted on the MoFlo ASTRIOS into 10% FBS RPMI-1640 media with 25 mM HEPES.

### RNA-seq analysis

RNA was isolated using Qiagen RNeasy Kit (Micro or Mini) or TRIzol. RNA integrity and concentration were determined using Eukaryote Total RNA Nano assay on BioAnalyzer. RNA-seq was performed on 10 ng–1μg of total RNA according to the Genewiz Illumina HiSeq protocol for poly(A)-selected samples (2 × 100 or 150 bp, ~350M raw paired-end reads per lane).

### MAJIQ splice junction analysis

The Modeling Alternative Junction Inclusion Quantification (MAJIQ) algorithm (v1.0.3a) was run with de novo junction detection, ten reads minimum per junction, each sample being quantified individually against the Ensembl hg38 GFF3 annotation (v94).

### Oxford Nanopore Technologies (ONT) long-read sequencing

Total RNA was extracted using the Qiagen RNeasy kit following manufacturer’s instructions. The mRNA was isolated from 100 μg of total RNA using Dynabeads mRNA DIRECT kit (Invitrogen) and subjected to direct RNA (SQK-RNA002, ONT) library preparation. Each library was loaded into a Spot-ON Flow Cell R9 version (FLO-MIN106D, ONT) and sequenced in a MinION Mk1B device (ONT) for 48 hours. Raw Fast5 files were converted to fastq with guppy, followed by alignment to the gencode version of hg38 (v30) using minimap2; the resulting bam file was visualized using the Integrative Genomics Viewer.

### Polysome profiling

Whole cell lysate was prepared by incubating 2 × 10^7^ REH cells in lysis buffer supplemented with 100 μg/ml cycloheximide. It was layered onto a 10-50% sucrose density gradient, subjected to centrifugation at 35,000 rpm for 3 h, and fractionated into 23 0.5-ml fractions. UV absorbance was monitored at 260 nm to record the polysome profile and RNA content. After qRT-PCR, the relative distribution of specific transcripts across the gradient was calculated using the formula 2^Ct (fraction 1-X)^× 100/sum.

### Immunoprecipitation (IP) with FLAG antibody-agarose conjugate

Anti-FLAG M2 Affinity Gel (Sigma A2220) beads were washed twice with lysis buffer containing 1 × Halt protease and phosphatase inhibitors (Thermo Fisher 78446, 100×). FLAG beads were then applied to cell lysates and incubated at 4°C under rotary agitation for 2 h. The mixtures were centrifuged at 1300 × g for 3 minutes at 4°C and washed with lysis buffer 3 times. To elute, samples were incubated in 0.1M Glycine HCl buffer (pH 2 −3) for 10 minutes at 4°C under agitation. Eluted supernatant was collected after centrifugation and neutralized by addition of 1M Tris-HCl (pH 8) equivalent to 40% final volume.

### SDS-PAGE and immunoblotting

Samples were mixed with reducing sample buffer (Thermo Fisher 39000, 5× containing DTT) and 12.5% final concentration of 2-Mercaptoethanol. To avoid aggregation of transmembrane proteins, samples were not boiled before running on 7.5% TGX gels (Biorad 4561023) at 130V. Samples were transferred to Immobilon-P Membrane (Millipore Sigma IPVH00010) at 100V for 90 minutes on ice. Membrane was blocked in 5% milk in TBST (0.1% Tween-20) at room temperature for 30 minutes before incubated with primary antibody at 4°C for 16 h, secondary antibody at room temperature for 30 minutes, and Immobilon Western Chemiluminescent HRP Substrate (Millipore WBKLS0500) for 5 minutes. Blots were visualized on Syngene G:Box Chemi XX6 imaging systems.

### Data-dependent acquisition (DDA) mass spectrometry (MS)

Samples were processed, cleaned-up, and Trypsin/LysC-digested prior to reverse phase HPLC and MS/MS. MS data were acquired on Thermo Fisher Q Exactive HF Orbitrap at the CHOP Proteomics Core.

### Morpholino (MO) transfection

Random control oligo and custom *FBXW7* MOs (Gene Tools) were electroporated into REH cells via the Neon transfection system (1325V, 10ms, 3 pulses) at 5 μM in 2-mL culture media. After 48 hours, cells were used for downstream assays.

### Mixed primers RT-PCR

Total RNA was reverse-transcribed using random primers and Multiscribe (Invitrogen). For PCR, all three *FBXW7* first coding exon forward primers were mixed with the reverse primer in coding exon 2. Platinum Taq (Invitrogen) was used for 30 cycles of amplification. PCR products were visualized following gel electrophoresis. Individual bands were gel-purified and Sanger sequenced.

### Taqman real-time qRT-PCR

qRT-PCR was performed using Taqman Master Mix (Applied Biosystems 4369016). Reactions were performed on Applied Biosystems QuantStudio 7 Flex and analyzed with the embedded software. Comparative CT was calculated against *GAPDH* expression within the sample. Relative quantity of all samples was then normalized to the control sample within the experiment.

### Cell cycle analysis

Cells were processed with FITC BrdU Flow Kit (BD 559619) for BrdU and 7-AAD staining following manufacturer’s instructions. Data were collected with FL1 and FL3 on Accuri C6.

### Cell death analysis

Cells were processed with FITC Annexin V Apoptosis Detection Kit (BD 556547) for Annexin V and PI staining following manufacturer’s instructions. Data were collected with FL1 and FL2 on Accuri C6. A dead cell positive staining control was prepared by MO control transfection, 16-h serum-free RPMI incubation, and flow cytometric staining.

## Results

### Normal human pro-B cells express FBXW7β as the predominant transcript isoform

To capture the extent of transcript variation in normal human B-cell development, we isolated hematopoietic progenitors and B-cell subsets from the bone marrow and tonsils for RNA-seq (Figure 1A). Using the Modeling Alternative Junction Inclusion Quantification (MAJIQ) algorithm^20^, we compared splice junctions and exon usage in the early progenitors, pro-B, pre-B, immature, and naïve mature B cells (Figure 1A). In these comparisons, thousands of genes were found to harbor at least one local splicing variation (LSV). Among them, 150 genes exhibit differential exon usage throughout the developmental process (Figure 1B). Gene Ontology (GO) analysis revealed that these 150 genes are overrepresented in biological pathways such as cell cycle process and positive regulation of proteolysis (Figure 1C). One of the genes, *F-box and WD Repeat Domain Containing 7 (FBXW7)*, is involved in four out of the top ten biological pathways identified in the GO analysis. FBXW7 is known as the substrate adaptor of the Skp1-Cullin-F-box containing (SCF) E3 ubiquitin ligase complex, and its three coding isoforms (α, γ, and β) are generated through distinct promoters and alternative first exon usage.^11^ While the α isoform is ubiquitously expressed, FBXW7β expression has not been reported in human B cells previously.

**Figure 1.**
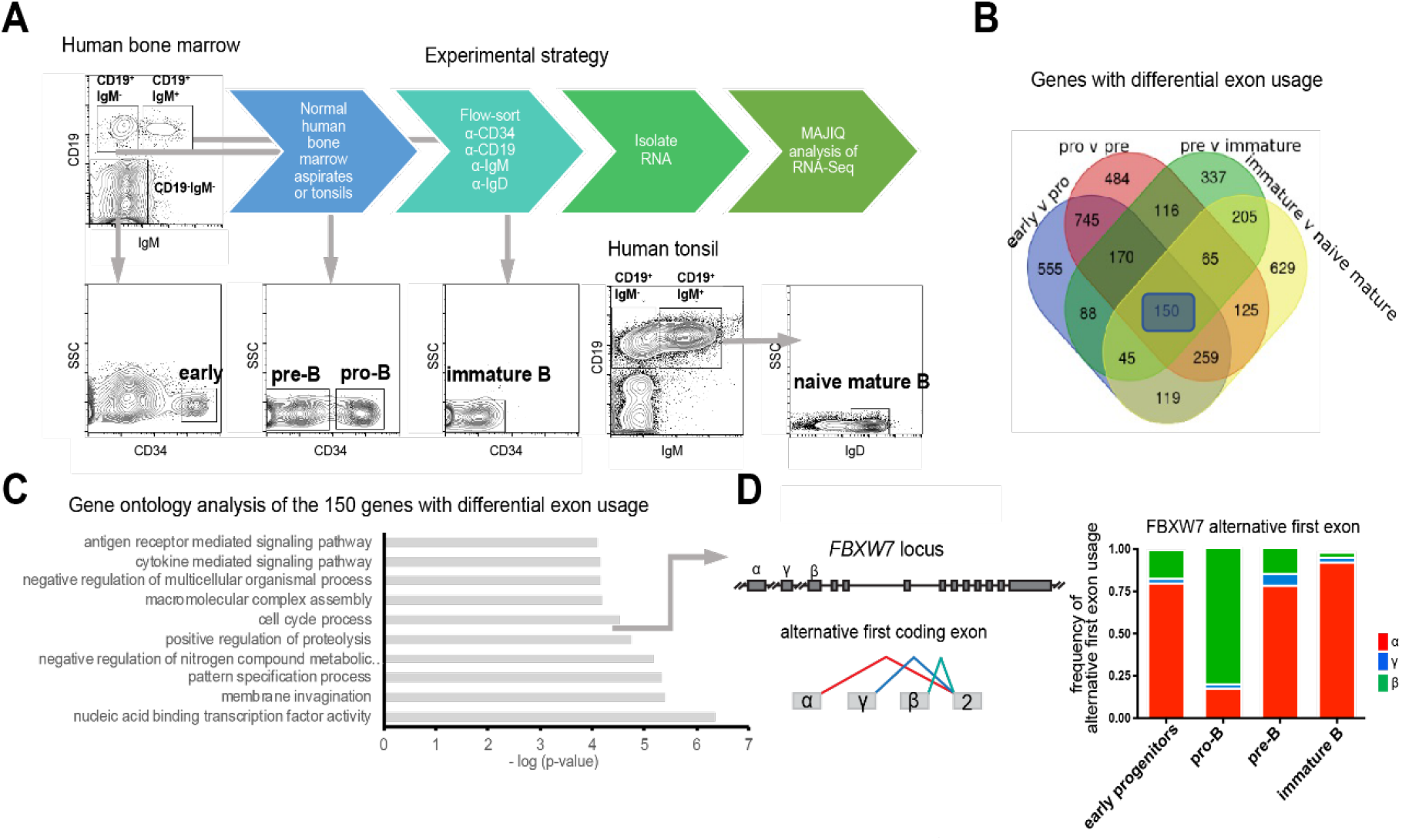
RNA-seq of primary normal human bone marrow and tonsil fractionations identifies *FBXW7* transcript variants. **(A)** Fluorescence-activated cell sorting and Modeling Alternative Junction Inclusion Quantification (MAJIQ) analysis of RNA-seq for normal human progenitors and B-cell subsets from the bone marrow (BM) and tonsils. **(B)** Venn Diagram of genes exhibiting differential exon usage throughout human B-cell development. Ensembl gene annotation was used. **(C)** Gene Ontology analysis of the 150 genes highlighted in (B). **(D)** Genomic configuration of *FBXW7* gene and its alternative first coding exon usage in normal BM subsets. Each stacked bar represents means of frequencies of α, γ, and β exon usage among 4 donors.

To our surprise, *FBXW7β* isoform appears as the predominant isoform in pro-B cells while the α isoform predominates in other stages of B-cell development (Figure 1D). The transition from early progenitors to pro-B is accompanied by the α-to-β isoform switch, which is reversed later (Figure 1D). We then investigated whether the Fbxw7 isoform switch occurs in other animal models. Upon analyzing existing transcriptional start site profiles (FANTOM5 Cap Analysis of Gene Expression) and RNA-seq data (NCBI GEO: GSE72018), we found no evidence of *Fbxw7β* expression in mouse hematopoietic tissues (Supplemental Figure 1).^21,22^ Thus, the FBXW7 α-to-β isoform switch may be unique to human B-cell development.

### Heterogeneous FBXW7 α and β transcript ratios in leukemic cell lines and primary B-cell Acute Lymphoblastic Leukemia (B-ALL) patient samples

After discovering *FBXW7β* expression in normal pro-B cells, we next sought to determine its expression status in hematologic malignancies. We visualized *FBXW7* coding isoform expression in a panel of leukemia cell lines from the Cancer Cell Line Encyclopedia (CCLE) (Figure 2A). Although Nalm6 and REH were both isolated from B-Cell Acute Lymphoblastic Leukemia (B-ALL) patients, Nalm6 expresses *FBXW7α* as the predominant isoform (highlighted in red), while REH predominantly expresses *FBXW7β* (highlighted in green in Figure 2A). We also examined primary B-ALL patient samples collected at the Children’s Hospital of Philadelphia (CHOP) and St. Jude Children’s Hospital (Figure 2B-C). B-ALL patients exhibit differential α-to-β ratios compared to normal counterparts (Figure 2B). We stratified patient samples based on disease subtypes and profiled their gene expression (Supplemental Figure 2). The Not Otherwise Specified, E2A, ETV6-RUNX1, and Philadelphia chromosome+ subtypes of B-ALL present heterogenous α-to-β transcript ratios (Figure 2C). However, *FBXW7α* predominates the ERG, hypodiploid, infantile, and MLL subtypes (Figure 2C). Intriguingly, usage of *FBXW7β* first exon is positively correlated with overall *FBXW7* transcript expression in hundreds of B-ALL patients (Figure 2D).

**Figure 2.**
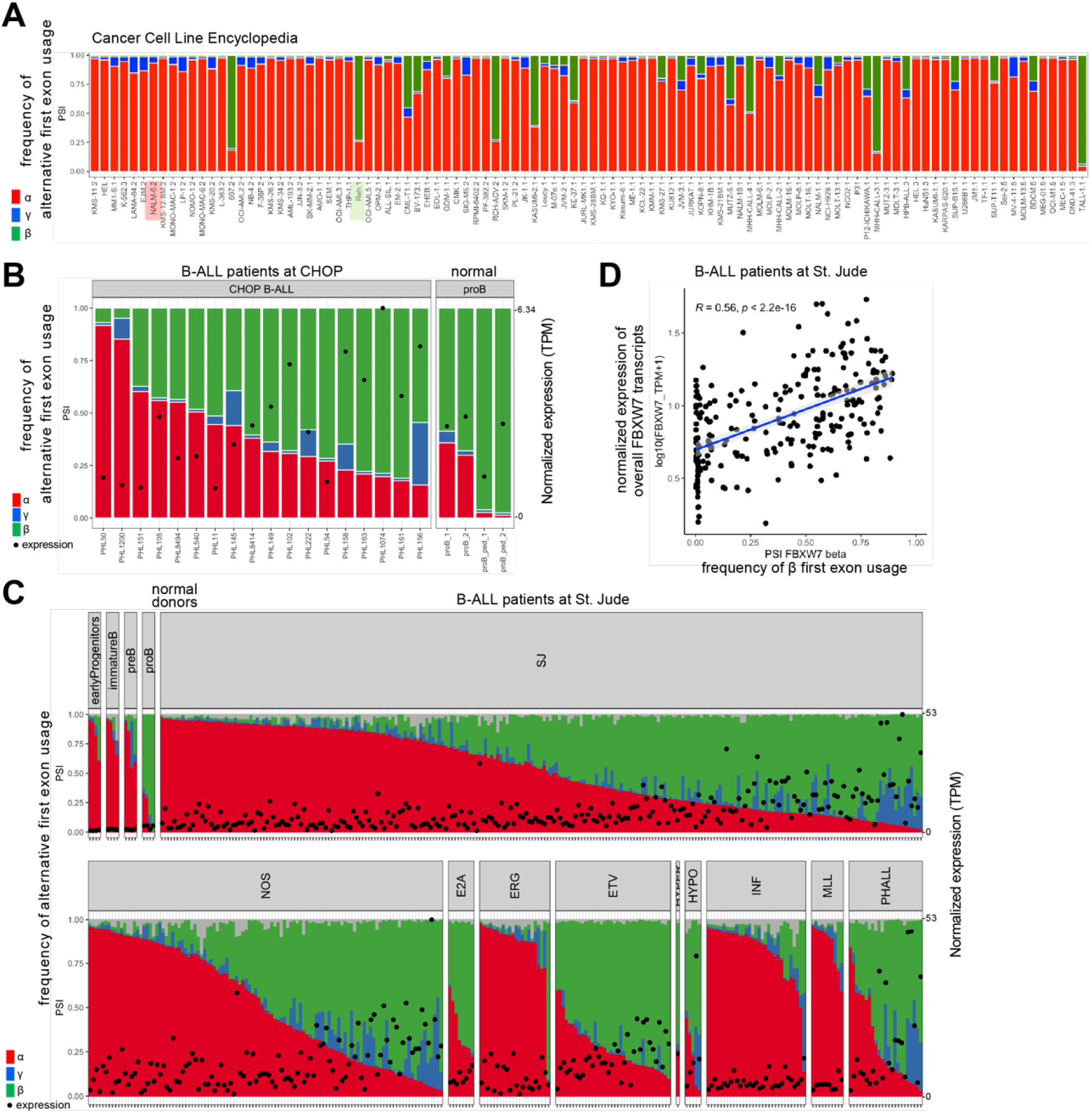
RNA-seq of leukemic cell lines and primary B-ALL patient samples revealed heterogenous *FBXW7 α* and *β* isoform ratios. **(A)** FBXW7 alternative first exon usage in leukemic cell lines from the Cancer Cell Line Encyclopedia RNA-seq repository. Nalm6 and REH B-Cell Acute Lymphoblastic Leukemia (B-ALL) cell lines are highlighted. **(B)** FBXW7 alternative first exon usage and transcript expression in B-ALL patients at the Children’s Hospital of Philadelphia (n=18) and normal BM donors (n=4). Each bar represents one sample. **(C)** FBXW7 alternative first exon usage and transcript expression in various B-ALL patient subtypes at St. Jude Children’s Hospital (n=247). **(D)** Spearman correlation of β exon usage and overall FBXW7 transcript expression in B-ALL patients at St. Jude Children’s Hospital.

### Full-length, translatable FBXW7β transcript is expressed in the REH B-ALL cell line and is enriched in the heavy polysome fraction

To further study the biology of *FBXW7* isoforms, we performed deep short-read RNA-seq on the REH cell line in-house. Consistent with our previous analysis of the CCLE data, *FBXW7β* stands as the predominant isoform in REH (Figure 3A). However, short-read technology only provides local information near the reference exon. To study co-occurring junctions in the same transcript, we used the Oxford Nanopore Technologies long-read RNA sequencing. Single-molecule, full-length *FBXW7β* transcript was successfully detected in the REH cell line (Figure 3B). Due to limitations of the long-read technology, the minor RNA isoforms in REH, *FBXW7α* and *γ*, were below the detection limit.

**Figure 3.**
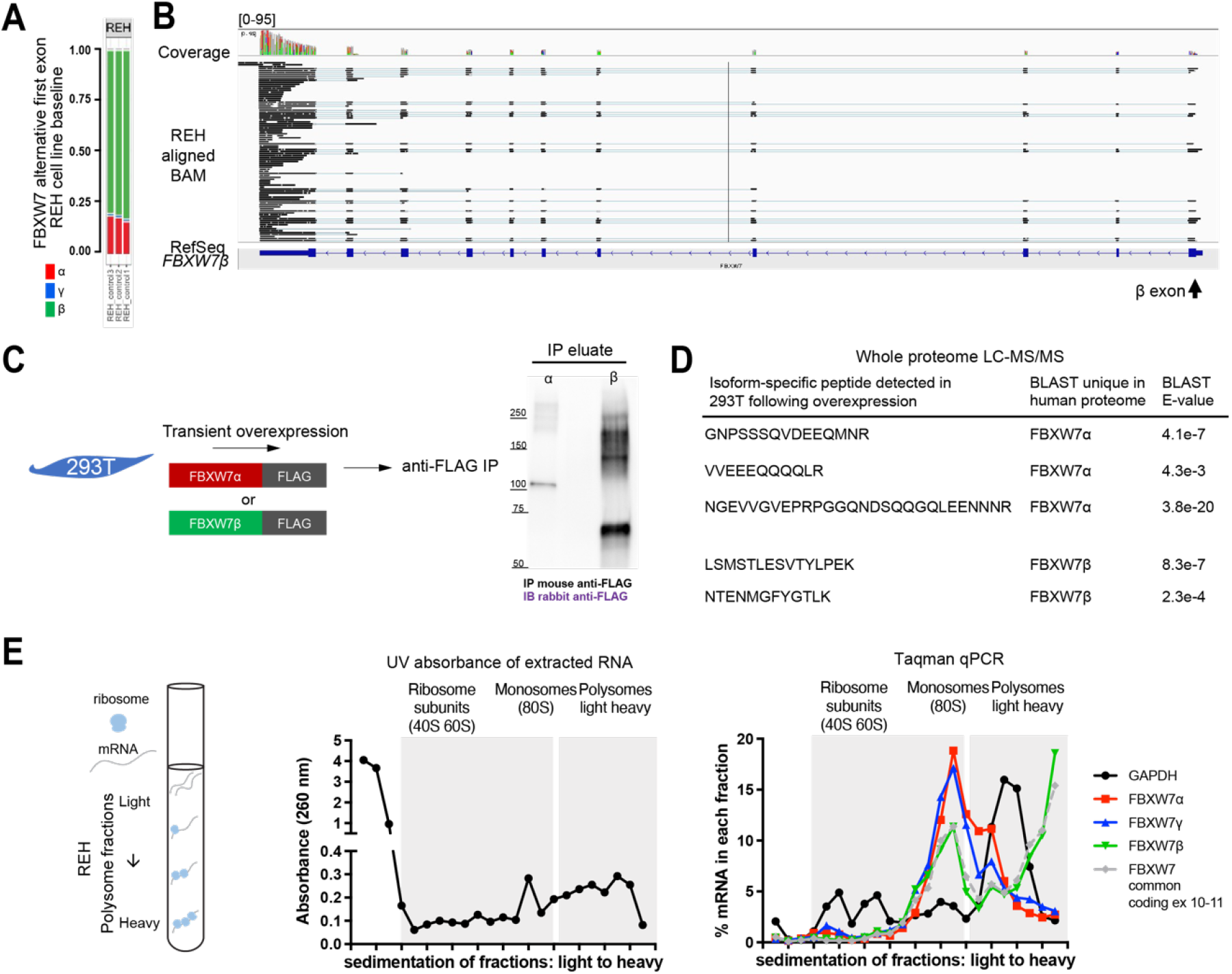
*FBXW7β* transcript is protein-coding and is enriched in the heavy polysome fraction. **(A)** MAJIQ analysis of FBXW7 alternative first exon usage based on short-read RNA-seq in REH. **(B)** Oxford Nanopore Technologies long-read RNA sequencing demonstrates full-length *FBXW7β* transcript expression in REH. **(C)** Detection of FBXW7 protein isoforms following robust transient overexpression in 293T cells, anti-FLAG immunoprecipitation (IP), and immunoblotting. **(D)** Data-Dependent Acquisition (DDA) mass spectrometry (MS) allows detection of overexpressed FBXW7 isoform-specific peptides in 293T cells, in addition to peptide sequences common to all isoforms. **(E)** Polysome profiling followed by qPCR. Specific mRNA distribution was expressed as a percentage of the total signal.

Detection of the FBXW7β protein remained challenging due to lack of suitable antibodies and short protein half-life.^23^ In our hands, all commercial antibodies failed to detect endogenous FBXW7β by immunoblotting (Supplemental Figure 3). Our subsequent attempts to generate mouse antibodies against FBXW7β-specific peptides did not lead to successful detection of FBXW7β (Supplemental Figure 4). To circumvent these technical limitations, we overexpressed FLAG-tagged FBXW7β in the 293T cell line for immunoprecipitation and detected the FBXW7β protein by immunoblotting (Figure 3C). Furthermore, we identified two independent β-specific peptides in the whole proteome by Data-Dependent Acquisition (DDA) mass spectrometry (Figure 3D). Besides isoform-specific peptides, peptides mapped to the regions common to all three coding isoforms were also found. To search for endogenous FBXW7 protein in the REH B-ALL cells, we next utilized the Data-Independent Acquisition (DIA) mass spectrometry. However, the DIA method failed to detect any FBXW7 peptides in the REH B-ALL cells. We thus investigated the translational status of the FBXW7 isoforms in these cells. To analyze mRNA distribution in relation to ribosome association, we performed polysome fractionation followed by qPCR. While *FBXW7α* and *γ* transcripts were found in the monosome fraction, *FBXW7β* transcript was predominantly found in the heavy polysome fraction in the REH cells (Figure 3E). This implied that *FBXW7β* transcript in REH cells is translated into a protein. To conclusively assign a function to that protein, we set out to perform both gain- and loss-of-function experiments in this cell line.

### Gain- and loss-of-function of FBXW7β isoform are well-tolerated in the REH B-ALL cell model

To discern relative contribution of each *FBXW7* coding isoform to cellular phenotypes, we generated a pan-*FBXW7* knockout (KO) single-cell clone and stably reconstituted individual isoforms (Figure 4A and Supplemental Figure 5). To this end, REH cells were first transduced with a Cas9 construct, yielding the REHCas9 cells. The REHCas9 cells were then transiently transfected with two CRISPR guide constructs simultaneously to remove the coding exon 2 of *FBXW7*, leading to a pan-*FBXW7* KO single-cell clone. Finally, the pan-*FBXW7* KO was stably transduced with individual FBXW7 isoform constructs that carried EGFP for Fluorescence-Activated Cell Sorting (FACS) selection. We encountered no difficulty expanding the transduced cultures, attesting to REH cells’ high tolerance for modest (6-40-fold) *FBXW7* overexpression (Figure 4B).

**Figure 4.**
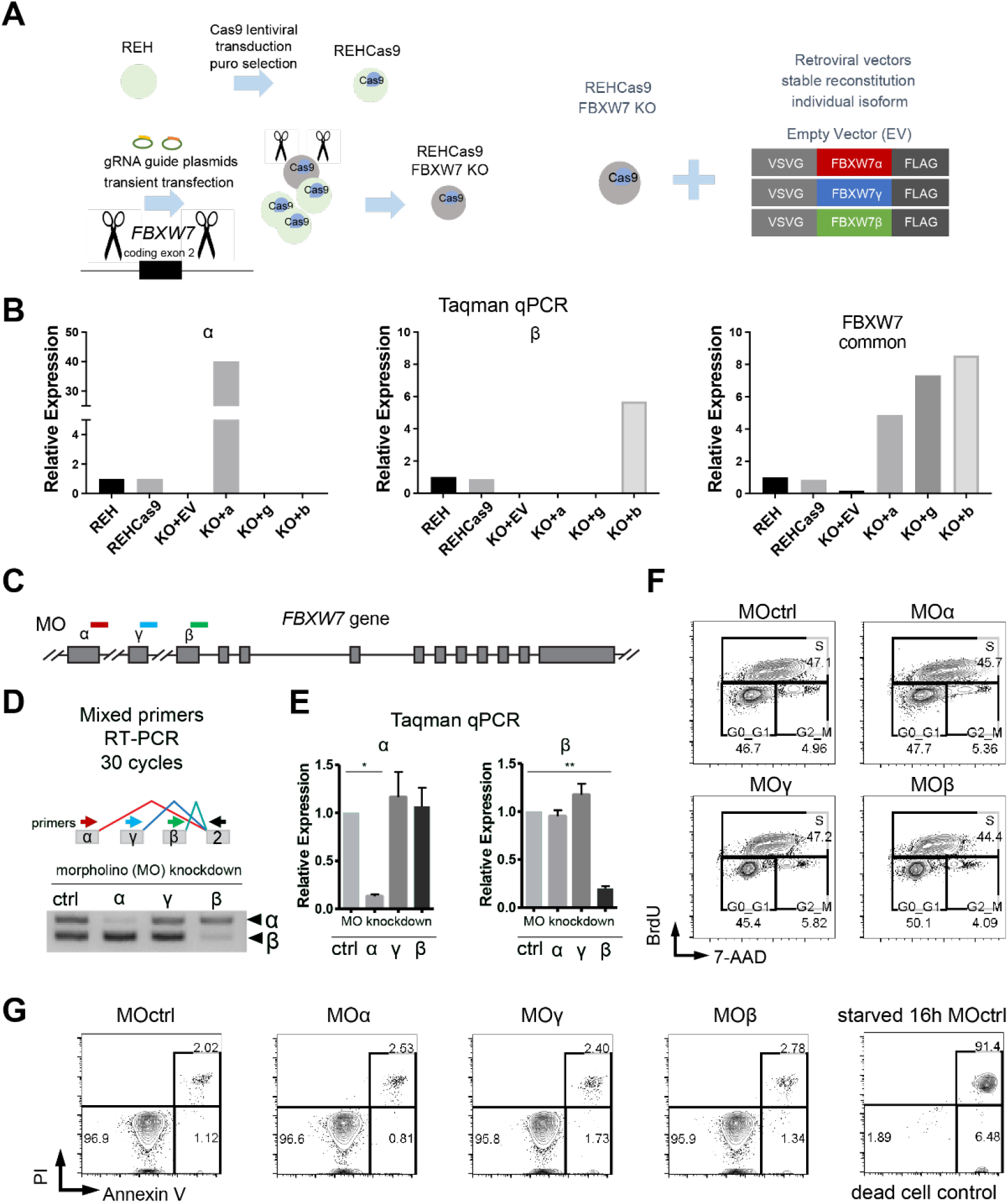
Generation of viable gain- and loss-of-function REH B-ALL cell models in which *FBXW7* isoforms were knocked out and reconstituted or transiently knocked down. **(A)** Experimental approach for stable reconstitution of *FBXW7* isoforms in a pan-*FBXW7* KO REHCas9 single cell clone. **(B)** Taqman qPCR to examine RNA expression in reconstituted REHCas9 *FBXW7* KO cells. Relative expression was normalized to REH parental cells. **(C)** Morpholinos (MO) were designed to target junctions of *FBXW7* alternative first coding exons and corresponding downstream introns for transient isoform knockdown (KD). **(D)** MO reduces FBXW7 transcripts in an isoform-specific manner. Mixed primer RT-PCR was performed following FBXW7 KD in REH cells. FBXW7α and β transcripts were detected while γ was below detection limit in the 30-cycle PCR. **(E)** Taqman qPCR of REH cells following FBXW7 isoform KD. Error bars represent means +/-SE of two independent experiments. One-way ANOVA of all groups were performed followed by Dunnett comparisons between the MO ctrl and other groups. \**Adj P* < .05, \*\**Adj P* < .01. **(F)** Transient FBXW7 isoform KD in REH B-ALL cells does not affect cell cycle progression. **(G)** Transient FBXW7 isoform KD in REH B-ALL cells does not affect cell viability. An independent MO ctrl treated sample was serum-starved overnight, acting as dead cell positive control. Data representative of two independent experiments in (D, F-G).

We also employed a loss-of-function approach via Morpholino (MO) splice site blocking (Figure 4C). The MO targets respective *FBXW7* first coding exon and its downstream intron, leading to transient isoform-specific knockdown (Figure 4D-E). In REH cells, *FBXW7* isoform ablation does not impact cell cycle progression or viability (Figure 4F-G). These two viable cell models, stable reconstitution and transient knockdown, facilitated our efforts to examine functions of *FBXW7β*.

### Transcriptome profiling identifies putative FBXW7β downstream targets in the two REH-based B-ALL cell models

As a member of the SCF E3 ubiquitin ligase complex, FBXW7 interacts with a protein substrate, leading to polyubiquitination of the substrate. The fate of the substrate depends on the type of ubiquitin linkage, such as the K48 and K63 polyubiquitin chains. The polyubiquitinated substrate may engage in proteasome-independent signal transduction or, more often, be degraded by the proteasome.^24^ This makes identification of direct FBXW7 targets challenging. To uncover secondary and tertiary effects of FBXW7α and β isoforms, we performed RNA-seq and profiled transcriptomes of the two REH-based B-ALL cell models (Figure 5A).

**Figure 5.**
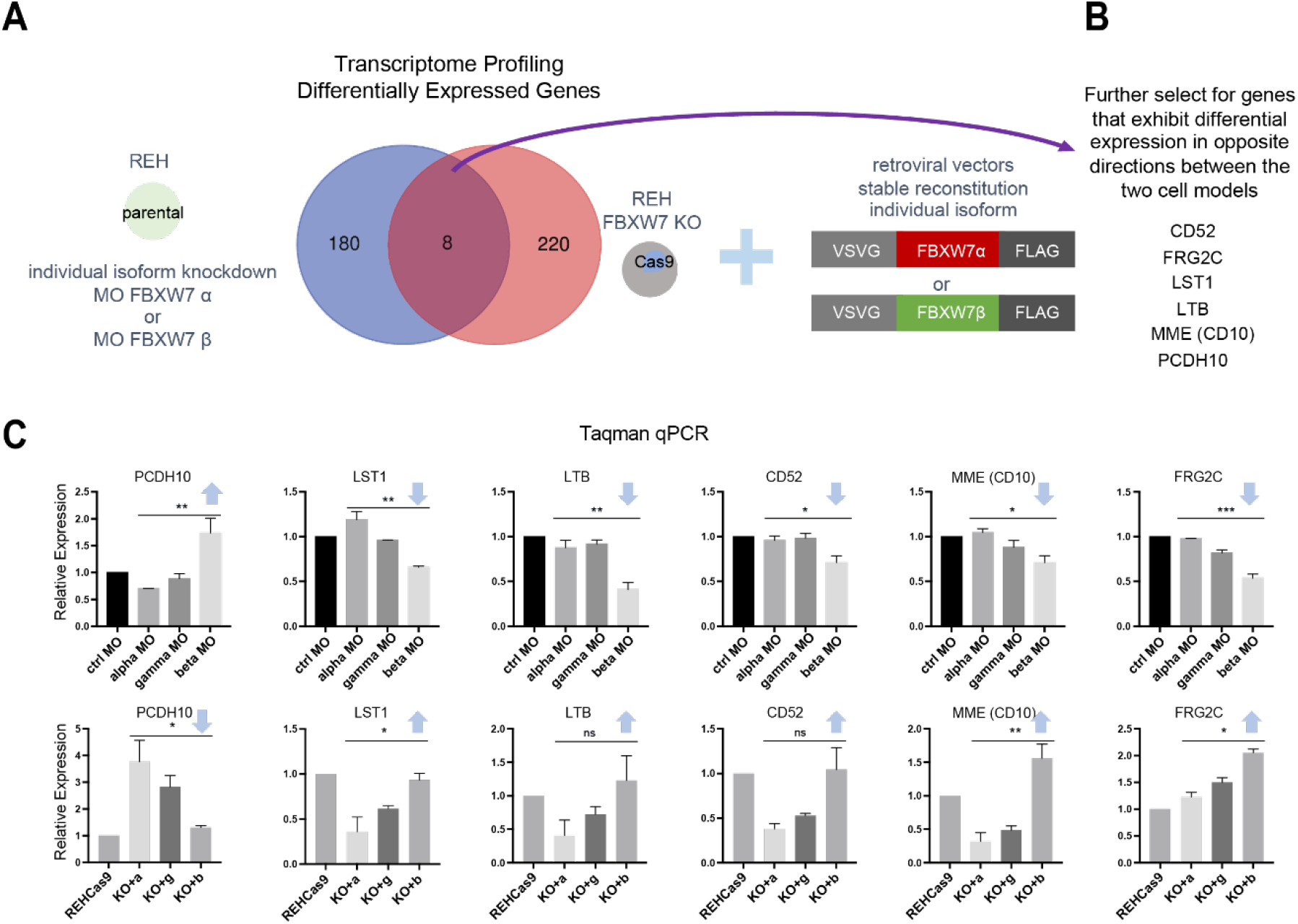
Transcriptome profiling of REH B-ALL cell models after *FBXW7* isoform knockdown or reconstitution reveals putative downstream targets. **(A)** Overlap of selected comparisons (α group v. β group) in RNA-seq data from the two REH cell models. Blue: Knockdown (KD) model gene list, differentially expressed genes in MO α v. MO β. Red: Reconstitution model gene list, differentially expressed genes in α reconstitution (KO+a) v. β reconstitution (KO+b). Each group includes replicates from two independent experiments. **(B)** List of overlapping downstream targets that exhibit opposite directionality of transcript foldchanges in the two models. **(C)** Taqman qPCR of the putative downstream target genes in the KD (top) and reconstitution (bottom) models. Arrows indicate directionality of transcript foldchanges comparing the α and β groups. Data represent means +/-SEM of two independent experiments. One-way ANOVA of all groups were performed followed by Tukey comparisons between the α and β groups. \**Adj P* < .05, \*\**Adj P* < .01, \*\*\**Adj P* < .001.

We first analyzed differential transcript expression between REH cells treated with MO *FBXW7α* or MO *FBXW7β*. We also compared transcript expression between pan-*FBXW7* KO REHCas9 cells reconstituted with either *FBXW7α* or *FBXW7β*. Approximately 200 differentially expressed genes were revealed in each comparison gene list. A subset of eight genes were found to overlap between the two lists (Figure 5A). In gain- and loss-of-function genetic perturbation models, putative gene targets are expected to be regulated in the opposite directions. Therefore, we further filtered the subset of genes based on directionality. As a result, six genes were differentially expressed in the opposite directions between the two models: *Protocadherin 10 (PCDH10), Lymphotoxin beta (LTB), Leukocyte specific transcript 1 (LST1), CD52, Membrane metalloendopeptidase (MME*, also known as CD10), and *FSHD region gene 2 family member C (FRG2C)* (Figure 5B). To corroborate the findings from the transcriptome analysis, we performed Taqman qPCR. Results demonstrated downregulation of *LTB, LST1, CD52, MME* (CD10), and *FRG2C* and upregulation of *PCDH10* in response to *FBXW7β* knockdown (Figure 5C top). In contrast, the opposite trend was observed in the reconstitution model, where effects of *FBXW7β* re-expression were compared to those of *FBXW7α* re-expression (Figure 5C bottom). These results suggest *FBXW7β* indirectly promotes *LTB, LST1, CD52, MME*, and *FRG2C* transcript expression.

### FBXW7β exon usage is positively correlated with the expression of the TNF superfamily and gene cluster in B-ALL patients

To determine whether these putative targets are associated with *FBXW7β* expression in leukemia samples, we next examined the large B-ALL patient cohort at the St. Jude Children’s Hospital (Figure 6A). Among the putative targets, the involvement of *LTB, LST1, CD52,* and *MME* (CD10) in inflammation has been reported. We found transcript expression of the four genes positively correlated to the frequency of *FBXW7β* exon usage (Figure 6A). As expected, there is no correlation between transcript expression of housekeeping genes (*EMC7, ACTB,* and *GAPDH*) and *FBXW7β* exon usage. On the other hand, *PCDH10* and *FRG2C* transcripts are barely expressed in the primary B-ALL patient samples.

**Figure 6.**
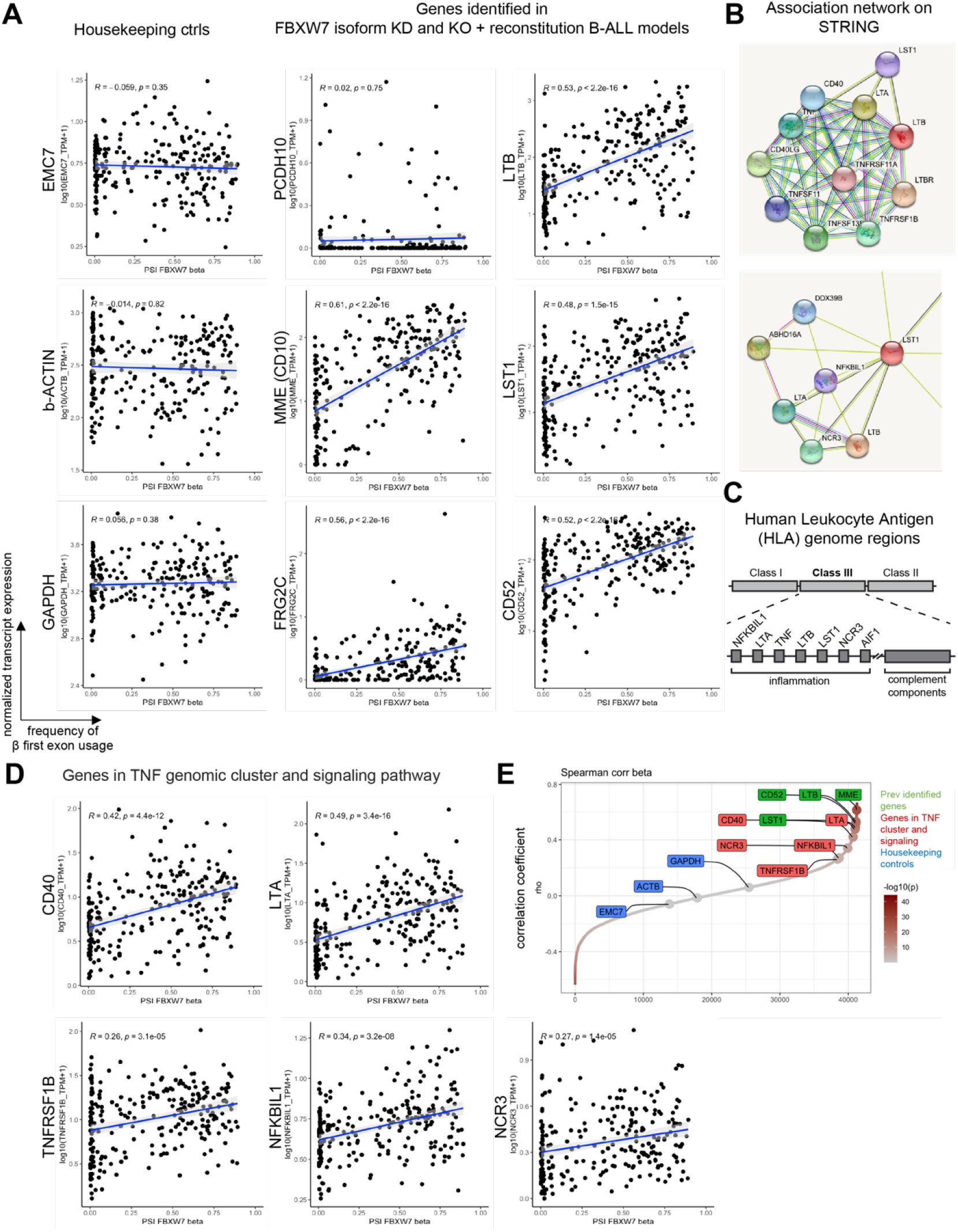
Transcript expression of *LTB, LST1, CD52, MME* (CD10), and several members in the *TNF* superfamily pathway is positively correlated with *FBXW7β* exon usage in BALL patients. **(A)** *FBXW7β* exon usage is positively correlated with *LTB, LST1, CD52,* and *MME* transcript expression in primary B-ALL patient samples at St. Jude Children’s Hospital. These four genes were identified in the FBXW7 isoform KD and KO + reconstitution REH B-ALL models (previous figure). **(B)** Top: STRING association network indicates functional association of *LTB* with members in the *TNF* superfamily signaling pathway. Bottom: STRING association network illustrates co-expression (black lines) of *LST1* with other genes. **(C)** Schematic of Human Leukocyte Antigen (HLA) genomic loci. Within the HLA Class III region, a gene cluster containing *TNF* is involved in inflammation. **(D)** Transcript expression of several members in the *TNF* superfamily signaling pathway is positively correlated with *FBXW7β* exon usage in B-ALL patients at St. Jude Children’s Hospital. Among those that exhibit positive correlation in (C) and (D), *LTB, LST1, NCR3, LTA*, and *NFKBIL1* gene loci are clustered in the same *TNF* genomic region. **(E)** Correlations of transcript expression from ~40,000 genes and the *FBXW7β* exon usage were analyzed. Genes were then ranked by the Spearman correlation coefficients (rho values). Positive rho values indicate positive correlations. Near zero rho values indicate lack of correlation.

To explore additional players that may interact with the putative targets, we turned to the STRING association network (Figure 6B).^25^ The STRING database consolidated prior knowledge from experiments, co-expression, and automated PubMed text mining. STRING indicated functional association of *LTB* with members in the *TNF* superfamily signaling pathway (Figure 6B top). *LST1* was associated with a network of genes based on co-expression prediction, including *LTB, LTA, NCR3,* and *NFKBIL1* (Figure 6B bottom). Interestingly, this network of genes is involved in inflammatory responses and housed in the same gene cluster containing *TNF* in the Human Leukocyte Antigen (HLA) Class III genomic region (Figure 6C).

Our analysis showed that transcript expression of members in the *TNF* superfamily signaling pathway and gene cluster (*CD40, LTA, LTB, TNFRSF1B, LST1, NCR3*, and *NFKBIL1*) positively correlates with *FBXW7β* exon usage in B-ALL patients (Figure 6A & D). Strength of correlations was also analyzed and ranked by the Spearman correlation coefficients (rho values) (Figure 6E). The expression of putative target genes identified in our two cell models (*LTB, LST1, CD52*, and *MME*) presented strong correlation with *FBXW7β* exon usage, while other members identified by the STRING functional association network presented moderate correlation. Taken together, our findings suggest a potential role of *FBXW7β* and its downstream effectors in inflammatory responses.

## Discussion

Most studies in malignancies, including T-ALL and colorectal cancers, identify *FBXW7* as a tumor suppressor.^26^ For example, recurrent missense mutations in *FBXW7* affect ubiquitination and half-life of the MYC oncoprotein, yielding leukemia-initiating cells.^27^ Paradoxically, *FBXW7* can also promote normal and malignant B-cell survival in various contexts.^8–10,28^ These reports highlight the pleiotropic functions of *FBXW7* that may be reprogrammed in response to cell-intrinsic and environmental cues. Regulation of *FBXW7* isoform expression adds another layer to its context-dependent roles. *FBXW7* encodes three coding isoforms (α, γ, and β) that are expressed from different promoters. These three protein isoforms differ in their N-termini but share the common nuclear localization signal (NLS), dimerization, F-box, and WD40 repeat domains. Of note, the α isoform contains an additional NLS, while the β isoform contains a unique transmembrane domain^11^,and the γ isoform is thought to be nucleolar.^29^

*FBXW7α* is the dominant isoform in most tissues, while *FBXW7β* is thought to be expressed in brain, and *FBXW7γ* in muscle. Most functional discoveries on FBXW7 are attributable to the α isoform. In colorectal cancer cells, FBXW7α carries out the bulk of the function.^23^ The FBXW7α isoform also enhances cell survival in the contexts of multiple myeloma and diffuse large B-cell lymphoma.^8,10^ However, studies that characterize the FBXW7β isoform are limited. Unexpectedly, our data showed *FBXW7β* as the dominant isoform in normal and malignant human pro-B cells. Our results provide evidence for full-length *FBXW7β* transcript expression and support its translational status in B-ALL cells, which is a necessary first step towards understanding the biological relevance of FBXW7β.

We found that usage of the *FBXW7β* exon is positively correlated with overall *FBXW7* transcript expression in primary B-ALL patient samples. This led to the question what allows precise regulation of FBXW7 isoform expression. Given that the germline promoter DNA sequences are identical across cell types that express distinct dominant FBXW7 isoforms, the difference in transcript expression may be attributed to transcription factor expression, epigenetic regulation, chromatin landscape, and/or post-transcriptional regulation mediated by microRNA. *FBXW7β* transcription is induced by p53.^30–32^ We did not find a correlation between *TP53* and *FBXW7β* transcript expression in the primary B-ALL patient samples (data not shown). Because post-translational modifications of p53 regulate its function as a transcriptional factor^33^, additional investigation is needed to clarify the involvement of p53 in the regulation of *FBXW7* in B-ALL. In addition, promoter hypermethylation has been suggested to inactivate *FBXW7β* expression in breast cancer.^18^ It remains to be determined whether promoter hypermethylation and other epigenetic mechanisms regulate *FBXW7β* expression in normal and malignant B-cell precursors.

We demonstrated functional consequences of *FBXW7β* isoform at the transcriptome level and uncovered *LTB, LST1, CD52*, and *MME* (CD10) as downstream targets. Among these recognized players in immunity, *LTB* and *LST1* reside in the tightly linked inflammatory *TNF* gene cluster in the HLA Class III genomic region.^34^ Furthermore, we found that members of *TNF* superfamily and gene cluster (*CD40, LTA, LTB, TNFRSF1B, LST1, NCR3*, and *NFKBIL1*) are associated with *FBXW7β* expression. The TNF superfamily controls expression of a constellation of genes through NFκB signal transduction and is critical in lymphoid development and regulation of immune responses.^35^ Our findings therefore establish a functional link to previous reports that implicate *FBXW7* in NFκB signaling and broaden the understanding of mechanisms by which *FBXW7* controls gene expression.^10,28,36,37^

Various FBXW7 isoforms, in particular α and β, may target distinct substrates for ubiquitination to achieve temporal regulation of B-cell development at homeostasis or may play a pathological role in neoplastic transformation. The FBXW7β-specific substrates in B-ALL still await discovery. From a prognostic and therapeutic perspective, further research is warranted to reveal the role of FBXW7β and its downstream effectors in leukemogenesis and cytotoxicity. Augmentation of lymphotoxin signaling is known to inhibit tumor growth in multiple xenograft models.^38,39^ Moreover, activation of the TNF superfamily may enhance anti-tumor efficacy of cell-based therapies. For instance, LTα has been shown to potentiate the T-cell response against melanoma and prolong survival in mice.^40^ Aiming to improve cell therapies, a recent study found that LTBR overexpression boosts T cell functions through activation of the canonical NFκB pathway.^41^ Conversely, upregulation of the NFκB signaling in leukemic cells may enhance cell survival and promote cancer progression. The balance between pro- and anti-tumor immune regulation is most likely shaped by factors intrinsic to cancer cells and the environmental stimuli.

## Acknowledgements

We thank former and present ATT lab members, former CHOP Clinical Hematopathology technical analyst Nicholas O’Grady, and CHOP ENT surgeons Steven Sobol and Steven Handler for their assistance with sample acquisition. We thank Wei Tong, Kristen W. Lynch, David Allman, Rizwan Saffie, Luca Busino, Steven H. Seeholzer, and Hua Ding for consultation on experimental design. We thank Junwei Shi, Krista Budinich, and Caroline R. Bartman for consultation and their LentiV_Cas9_puro (cc60) and Lenti_gRNA-GFP(LRG)_2.1T CRISPR plasmids as generous gifts.

This work was supported by the grants from the National Institutes of Health (U01 CA232563 to YB and ATT, T32 CA115299 and TL1 TR001880 to SYY), St. Baldrick’s-Stand Up to Cancer (SU2C-AACR-DT-27-17 to ATT), the V Foundation for Cancer Research (T2018-014 to ATT) and by the Alex’s Lemonade Stand Foundation Innovation Award to ATT.

## Author contributions and Disclosures

SYY designed research, performed research, analyzed data, wrote the paper

KEH contributed vital new reagents or analytical tools, analyzed data

HF performed research, analyzed data

LAS performed research

MA performed research

KLB performed research

ASN analyzed data

VP contributed vital new reagents

YB contributed vital analytical tools

KSJEJ designed research

ATT designed research, analyzed data, wrote the paper

The authors declare no relevant conflicts of interests.

## Notes

### Competing Interest Statement

The authors have declared no competing interest.

