## Supplementary material for "FBXW7β isoform drives transcriptional activation of a proinflammatory TNF cluster in normal and malignant pro-B cells": Supplem Methods and Figures

Yang *et al.*

Contains Supplemental Methods, Supplemental Tables, and Supplemental Figures

### Supplemental Methods

#### Dataset usage

We accessed RNA-seq data from the St. Jude Cloud, an initiative of St. Jude Children's Research Hospital (https://www.stjude.cloud) and from the Cancer Cell Line Encyclopedia (CCLE) (https://sites.broadinstitute.org/ccle). St. Jude Cloud data used for this analysis (EGAD00001002704 and EGAD00001002692) were accessed by permission from the Computational Biology Committee through the European Bioinformatics Institute (EMBL-EBI).

#### Spearman correlation

Correlations and their significance were computed using the nonparametric Spearman's rank-order correlation implemented in R function cor.test().

#### Cell culture

REH cells were maintained in RPMI-1640 medium supplemented with 10% FBS, 2 mmol/L l-glutamine, 25 mM HEPES, and antibiotic-antimycotic at 37°C and 5% $CO_2$. 293T cells were cultured in DMEM with the same supplements.

#### Cell lysis for immunoprecipitation (IP) and immunoblotting (IB)

Cells were centrifuged at 250 × g for 5 minutes. The cell pellet was lysed on ice in buffer containing 150mM NaCl, 50mM Tris pH 8.0, 1% Triton X-100, and 2 × Halt protease and phosphatase inhibitors (Thermo Fisher 78446, 100×). Protein concentration of lysates was maintained at 2 – 4 μg/μL. Cell debris was pelleted at 5000 × g for 5 minutes. Supernatant was collected for direct IB or IP before IB.

#### CRISPR-Cas9 genome editing and reconstitution

REH cells were stably transduced with LentiV_Cas9_puro (cc60) viral construct, yielding the REHCas9 cells. To generate a pan-FBXW7 KO single-cell clone, REHCas9 cells were transiently transfected with two CRISPR Lenti_gRNA-GFP(LRG)_2.1T DNA constructs simultaneously to remove the coding exon 2 of FBXW7. Finally, the pan-FBXW7 KO cells were stably reconstituted with individual FBXW7 isoform MIGR1 retroviral constructs that carried EGFP for Fluorescence-Activated Cell Sorting (FACS) selection.

#### Custom antibody generation

FBXW7β peptide-KLH conjugate antigen preparation, immunization of BALB/c and C57BL/6 mice, and antiserum preparation were performed by GenScript. Antigen peptide sequences are indicated in Supplemental Figure 4.

#### MG132 treatment

REH cells were left untreated or treated with 1:1000 DMSO vehicle or 10 μM MG132 (Peptide Institute 3175-v) for 4 hours in complete RPMI-1640.

### Supplemental Tables

| Supplemental Table 1. Taqman real-time qPCR primers and probes | | |
|---|---|---|
| Target | Type | Sequence 5'->3' |
| FBXW7al | Primer 1 | CGA ACT CCA GTA GTA TTG TGG ACC T |
| | Primer 2 | TTC TTT TCA TTT TTG TTG TTT TTG TAT AGA |
| | PrimeTime Probe | /56-FAM/CC CGT TCA C/ZEN/C AAC TCT CCT CCC CA/3IABkFQ/ |
| FBXW7ga | Primer 1 | GAG CCT CTA CCA CAT CAA ACT |
| | Primer 2 | AAA GAG CGG ACC TCA GAA CC |
| | PrimeTime Probe | /56-FAM/AG CAT TAG C/ZEN/A TCA TTG CCC AAG GC/3IABkFQ/ |
| FBXW7be | Primer 1 | CAG CCG GAC ACA CGG G |
| | Primer 2 | GGT CCA ACT TTC TTT TCA TTT TGT AAA |
| | PrimeTime Probe | /56-FAM/AA ATA CAG A/ZEN/A AAT ATG GGT TTC TAC GGC ACA TTA AAA ATG /3IABkFQ/ |
| FBXW7common(ex10-11) | Primer 1 | TGG GAT ATC AAA ACA GGA CAG TGT |
| | Primer 2 | TAA ACA GGT CAC AGC ACT CTG ATG |
| | PrimeTime Probe | /56-FAM/ACAAACATT/ZEN/GCAAGGTCCCAACAAG/3IABkFQ/ |
| GAPDH | Primer 1 | TGT AGT TGA GGT CAA TGA AGG G |
| | Primer 2 | ACA TCG CTC AGA CAC CAT G |
| | PrimeTime Probe | /56-FAM/AA GGT CGG A/Zen/G TCA ACG GAT TTG GTC /3IABkFQ/ |
| CD52 | Primer 1 | GTT TGG CTG GTG TCG TTT |
| | Primer 2 | GCC ACG AAG ATC CTA CCA AA |
| | PrimeTime Probe | /56-FAM/TGA GAG TCC /ZEN/AGT TTG TAT CTG TAC CAT AAC CA/3IABkFQ/ |
| LST1 | Primer 1 | GTC CCT GAT CCC TGA CCT AA |
| | Primer 2 | GAA GGA CCA CTG CCA GAA G |
| | PrimeTime Probe | /56-FAM/TCG CGG AAT /ZEN/GAT GAT ATA TGT ATC TAC GGG /3IABkFQ/ |
| LTB | Primer 1 | CAG CTG CCC ACC TCA TAG |
| | Primer 2 | ACA GTA GAG GTA ATA GAG GCC G |
| | PrimeTime Probe | /56-FAM/AAA CGC CTG /ZEN/TTC CTT CGT CGT CT/3IABkFQ/ |
| MME | Primer 1 | AAG AAA CAG CGA TGG ACT CC |
| | Primer 2 | GCA GCT GAT TTT ATG CAG TCT G |
| | PrimeTime Probe | /56-FAM/AGG AGC AGG /ZEN/ACA AGG ACC GAG A/3IABkFQ/ |
| FRG2C | Primer 1 | TTC CAA GGA TAT CTG CCA AGA C |
| | Primer 2 | CAG CAG TGG AGG ATC TTG ATT |
| | PrimeTime Probe | /56-FAM/CCA GAA GAG /ZEN/GAG TGC AAC TTG ACG T/3IABkFQ/ |
| PCDH10 | Primer 1 | CTC AGT GCA ATT GGA GAA GAG A |
| | Primer 2 | CCA GGA AGC CGA CAT AGT AAG |
| | PrimeTime Probe | /56-FAM/AGT GGT CAT /ZEN/GGA GAC AGT GAA CAG G/3IABkFQ/ |

| **Supplemental Table 2. Regular PCR primers** | |
|---|---|
| RT-PCR mixed primers | Sequence 5'->3' |
| FBXW7_syy_al_F | CAGCAAAAGACGACGAACTGG |
| FBXW7_syy_ga_F | CAGGACATTTGGTAGGGGAAGG |
| FBXW7_syy_be_F | TGACAGGGCATAGTCTCCTCC |
| FBXW7_syy_2_R | AAAGAGCGGACCTCAGAACC |
| Genotyping PCR primers flanking coding exon 2 | Sequence 5'->3' |
| FBXW7_syy_gD2_F | AGCCTAATAACTGTGAGAGTGGG |
| FBXW7_syy_gD2_R | AAGGGAAGAAACCAGCCAGATC |

| **Supplemental Table 3. CRISPR guide and Morpholino sequences** | | |
|---|---|---|
| CRISPR guide combination for coding exon 2 removal | | |
| CRISPR guide | Sequence 5'->3' | |
| all FBXW7 sg#1 | CTTACGACATTAGGGGCTAG | |
| all FBXW7 sg#2 | CTAGGGTAGACATTTATGTA | |
| Morpholinos | | |
| Target | Name | Sequence 5'->3' |
| FBXW7al | MOalpha | ATTGAATATACTCACTTTTGTTGTT |
| FBXW7ga | MOgamma | TTCAAATGTGTGAGACTTACCCGTC |
| FBXW7be | MObeta | GAAGAAAACAGCTTACTTACTTTGT |
| Random control oligo | MOctrl | 25N, mixture of up to 4^25 different sequences |

| **Supplemental Table 4. Flow cytometry and immunoblot reagents** | | | |
|---|---|---|---|
| Flow cytometry reagents | | | |
| Specificity | Fluorochrome | Manufacturer | Catalog Number |
| Human TruStain FcX | | Biolegend | 422302 |
| CD34 | PE | Beckman Coulter | IM1459U |
| IgM | FITC | Beckman Coulter | B30655 |
| CD19 | APC | Beckman Coulter | IM2470U |
| IgD | PE | Thermo Fisher | 12-9868 |
| Immunoblot antibodies | | | |
| Specificity | Manufacturer | Catalog Number | |
| FLAG | Cell Signaling Technology | 2368S | |
| FLAG | Sigma | F7425 | |
| FBXW7 | Abcam | ab109617 | |
| FBXW7 | Thermo Fisher | 40-1500 | |
| FBXW7α | Bethyl | A301-720A | |
| β-ACTIN HRP | Cell Signaling Technology | 12262 | |
| GAPDH HRP | Cell Signaling Technology | 3683S | |
| anti-mouse IgG HRP | Cytiva | NA931-1ML | |
| anti-rabbit IgG HRP | Cytiva | NA934-1ML | |

### Supplemental Figures

A

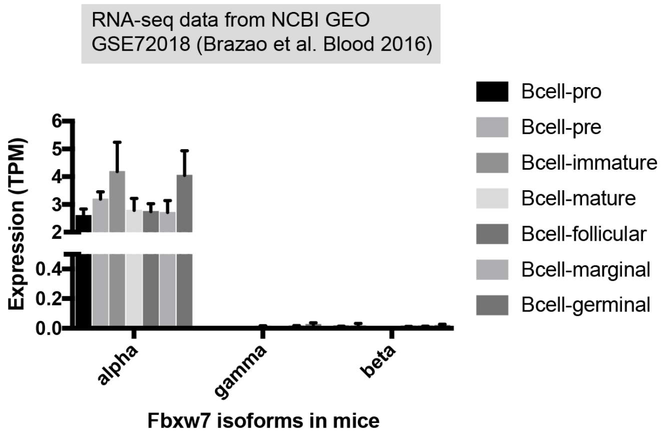


B

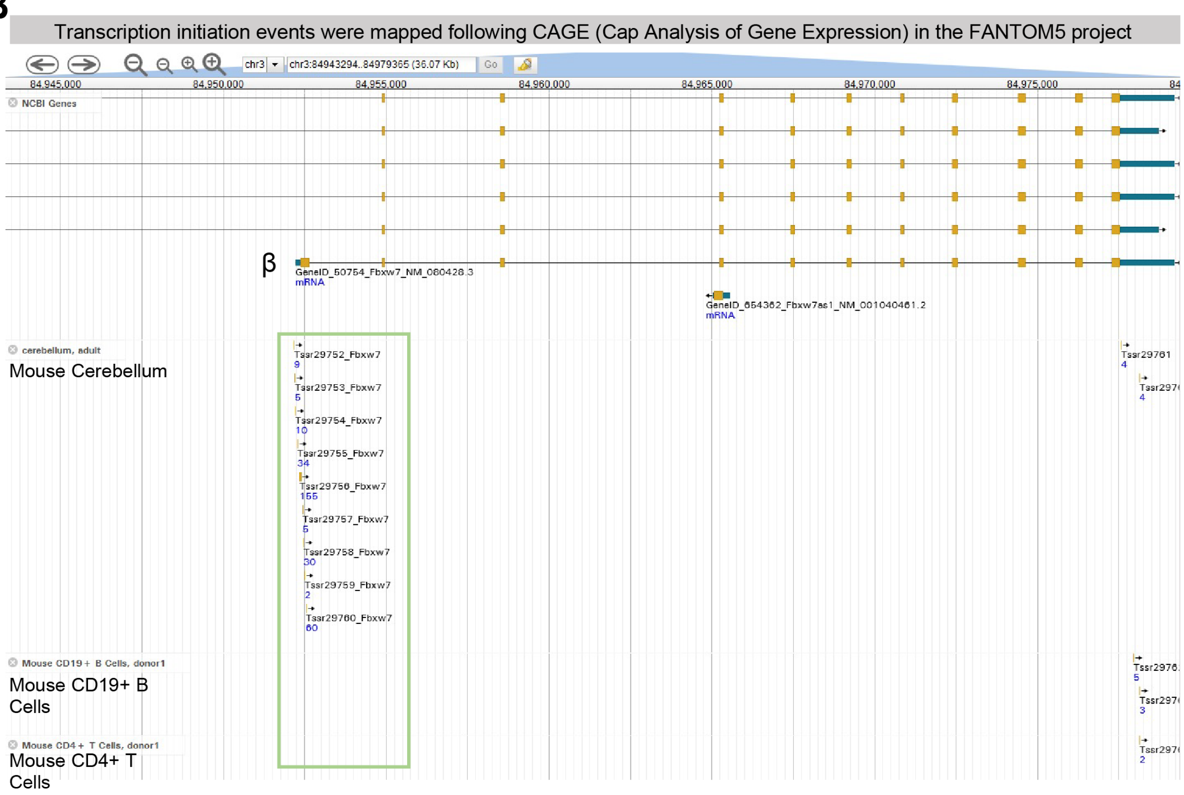


**Supplemental Figure 1. Existing RNA-seq and 5' Cap Analysis of Gene Expression (CAGE) datasets provide no evidence of *Fbxw7β* expression in mouse hematopoietic tissues. (A)** RNA-seq of various mouse B-cell subsets from NCBI GEO GSE72018 (Brazao T et al. Blood 2016) was reanalyzed to quantify *Fbxw7* isoform expression. HTSeq v0.8.0 was used in intersection-strict mode against the Ensembl mouse gene annotation (GRCm38/mm10 v87). **(B)** FANTOM5 CAGE transcriptional start site mapping dataset. Mouse *Fbxw7β* RNA expression is detected in nervous tissues but not in lymphocytes. Viewed on Mouse Genome Informatics (MGI) at JAX http://jbrowse.informatics.jax.org/?data=data%2F-mouse&loc=chr3%3A84943294..84979365&tracks=DNA%2CENSEMBL%2CNCBI%2C15-8B2%2C11856-125A2%2C11854-124I9&highlight=

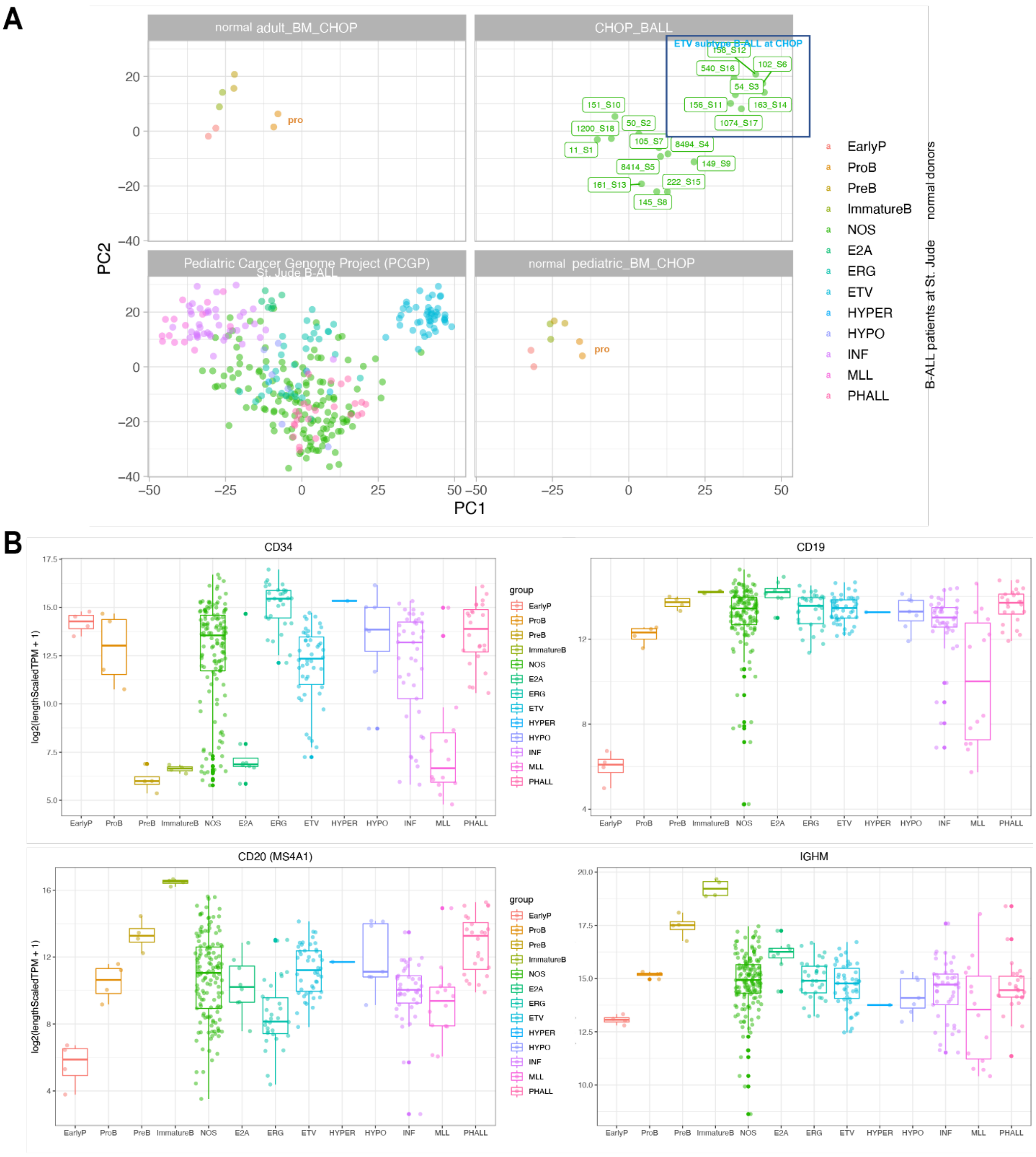


normal donors B-ALL patients at St. Jude

**Supplemental Figure 2. Gene expression of primary B-ALL patient samples compared to normal counterparts. (A)** Principal component analysis of transcriptome profiles of B-ALL patients at Children's Hospital of Philadelphia (CHOP) (n=18), St. Jude Children's Hospital (n=313), and normal BM cell subsets. **(B)** Transcript expression of B-cell and hematopoietic markers in B-ALL patients at St. Jude Children's Hospital (n=313) and in normal BM cell subsets (early progenitors, pro-, pre-, immature B cells). Salmon algorithm (https://combine-lab.github.io/salmon), tximport, DESeq2, and ggplot2 R packages were used for analysis.

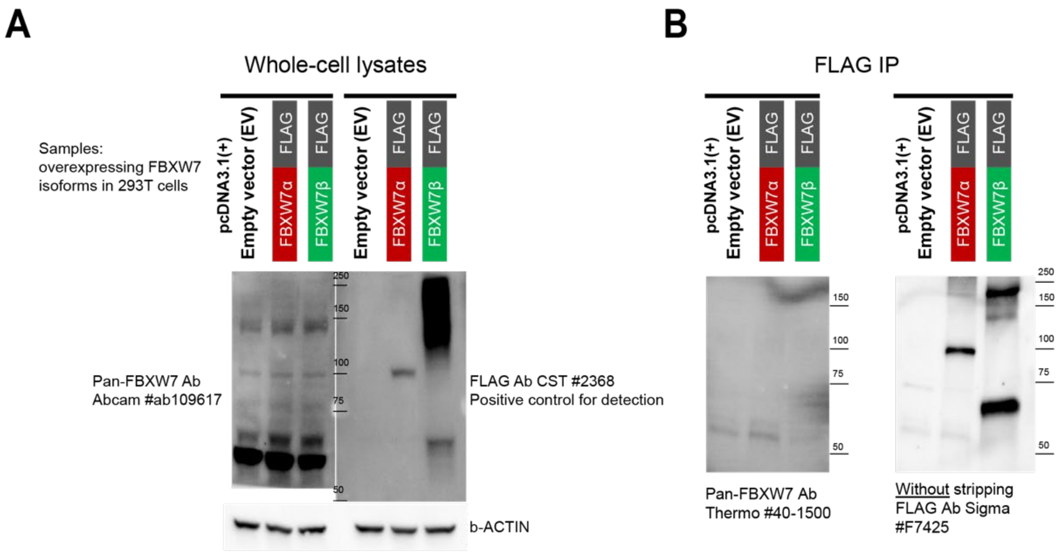


**Supplemental Figure 3. Commercial antibodies against pan-FBXW7 epitopes fail to detect FBXW7. (A)** Western blot of 293T cells transiently transfected with *FBXW7* isoform overexpression constructs using various antibodies. **(B)** 293T cells were transiently transfected with *FBXW7* isoform overexpression constructs, lysed, immunoprecipitated with anti-FLAG agarose beads, and immunoblotted with Thermo #40-1500 FBXW7 antibody first. Without stripping, the same membrane was blotted with FLAG antibody Sigma #F7425. Overexpression of FBXW7 protein isoforms was confirmed with the FLAG antibody.

**A**

Peptide sequence encoded by beta exon
**Bold**: peptide antigen for raising custom mouse antisera

1st attempt: MCVPRSGLILSCICLYCGVLLPVLLPNLPFLTCLSMSTLESVTYLPEKGLYCQRLPSSR**THGGTESLKGKNTE**NMGFYGTLKMIFYK
2nd attempt: MCVPRSGLILSCICLYCGVLLPVLLPNLPFLT**CLSMSTLESVTYLPEKGLY**CQRLPSSRTHGGTESLKGKNTENMGFYGTLKMIFYK

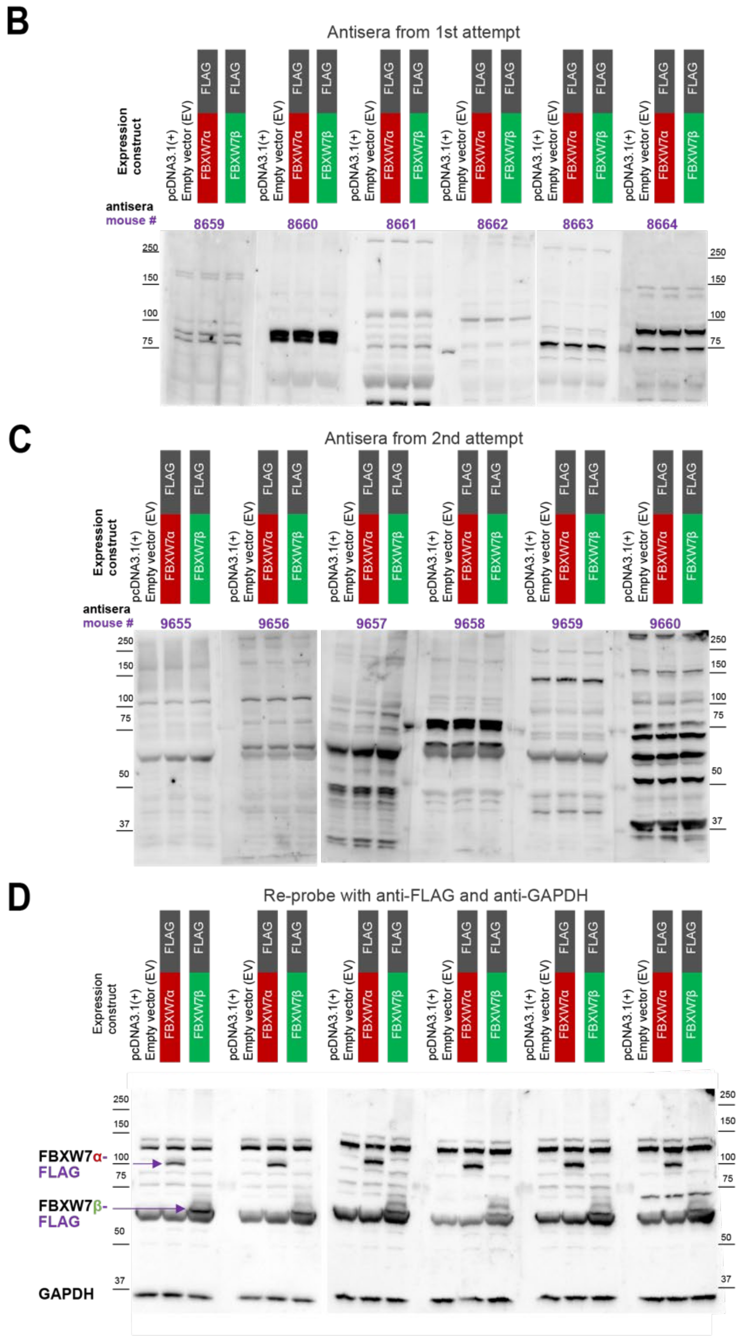


**Supplemental Figure 4. Attempts to generate FBXW7β-specific antibodies did not lead to successful detection. (A)** Amino acid sequence encoded by the FBXW7 β exon. Highlighted sequences were used as two different immunogens for two antisera generation attempts. **(B)(C)** Antisera Western blot compared to **(D)** anti-FLAG Western blot in 293T cells transiently transfected with *FBXW7* isoform overexpression constructs. Presence of overexpressed isoforms was confirmed after stripping anti-FBXW7β antisera and re-probing with anti-FLAG (Sigma #F7425).

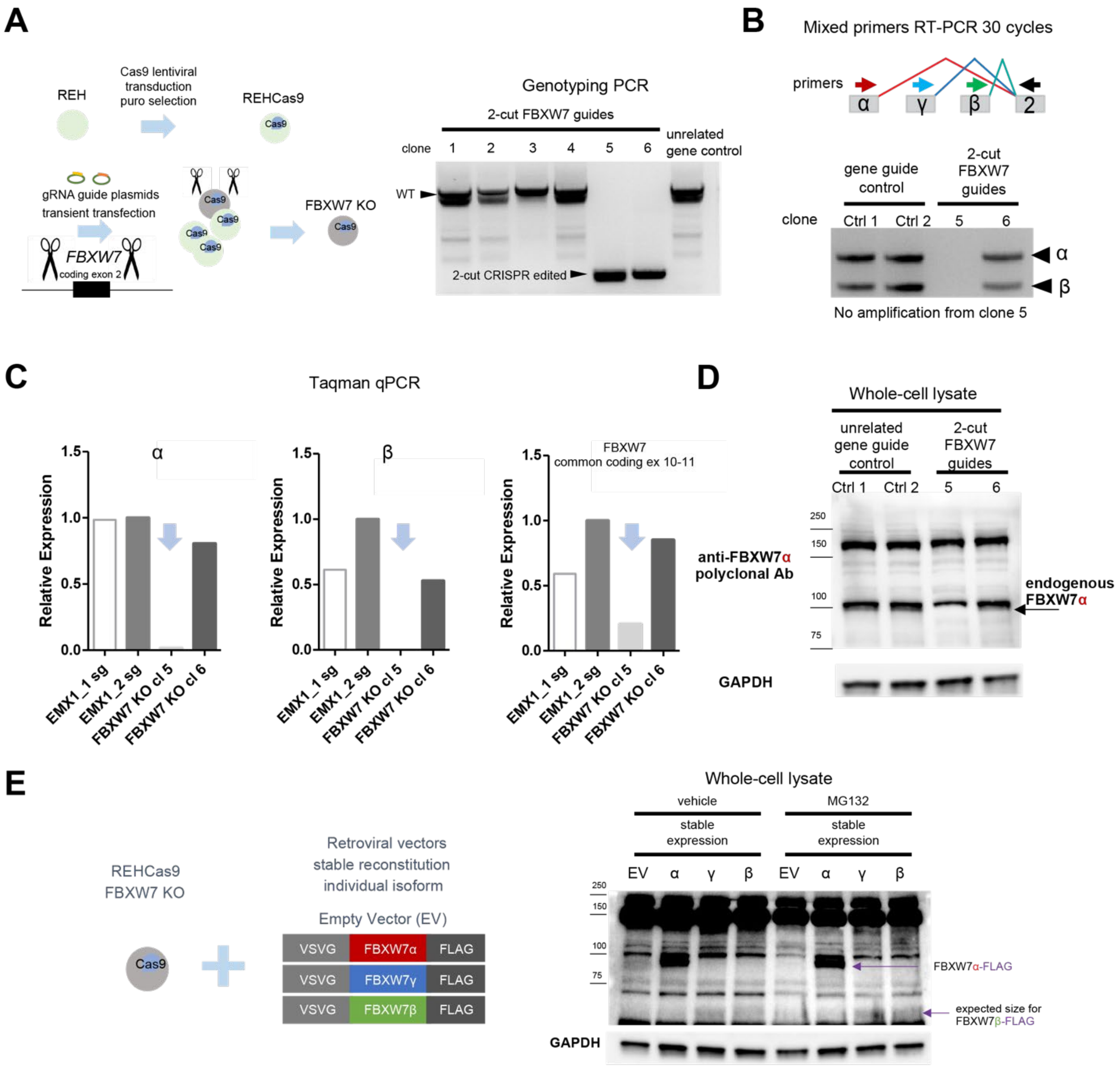


**Supplemental Figure 5. Generation of pan-*FBXW7* KO REHCas9 single cell clone with 2-cut CRISPR-Cas9 genome editing followed by reconstitution. (A)** Genotyping KO single cell clones with genomic DNA PCR: short amplicon reflects removal of *FBXW7* coding exon 2 shared by all isoforms. The control sample was transfected with one CRISPR guide targeting an unrelated gene *EMX1*. **(B)** Mixed primers RT-PCR (30 cycles) to examine RNA expression of KO single cell clone after 2-cut CRISPR-Cas9 genome editing. **(C)** Taqman qPCR to examine RNA expression of KO single cell clone after 2-cut CRISPR-Cas9 genome editing. Transcript quantity was relative to control sample EMX1_2 sg, where one CRISPR guide was used to target the unrelated gene *EMX1*. **(D)** Lack of FBXW7α protein expression in KO single cell clone, Western Blot with anti-FBXW7α (Bethyl rabbit polyclonal #A301-720A). **(E)** Stable reconstitution of REHCas9 FBXW7 KO cells followed by Western Blot with anti-FLAG (CST #2368). FBXW7 α-FLAG protein is readily detected. FBXW7β-FLAG is detected at ~64 kDa. MG132 treatment does not affect FBXW7 protein levels. Results representative of 3 independent experiments.
